# Virion stripping: A new role for bacterial proteases

**DOI:** 10.1101/2025.05.03.652018

**Authors:** Carlos J. Rodriguez Hernandez, Marina Terekhova, Phylip Chen, John Perpich, Barbara Potempa, Katherine Carey, Mark E. Peeples, Mitchell H. Grayson, Jan Potempa, Maxim Artyomov, Richard J Lamont, Juhi Bagaitkar

**Affiliations:** Center for Microbe and Immunity Research, The Abigail Wexner Research Institute at Nationwide Children’s Hospital, Columbus, OH, USA; Department of Pathology and Immunology, Washington University School of Medicine, Saint Louis, MO 63110, USA; Department of Pharmaceutical Sciences, College of Pharmacy and Health Sciences, Sullivan University, Louisville, KY, USA; Department of Oral Immunology and Infectious Diseases, School of Dentistry, University of Louisville, Louisville, KY, USA; Center for Clinical and Translational Research, The Abigail Wexner Research Institute at Nationwide Children’s Hospital, Columbus, OH, USA; Division of Allergy and Immunology, Department of Pediatrics, Nationwide Children’s Hospital and The Ohio State University College of Medicine, Columbus, OH, USA; Department of Pediatrics, The Ohio State University College of Medicine, Columbus, OH, USA

## Abstract

We demonstrate that proteases produced by the oro-pharyngeal bacterial colonizer *Porphyromonas gingivalis (Pg)* reduce viral burden and modulate host interferon responses during respiratory syncytial virus (RSV) infection. Several oral bacteria, including *Pg*, have been shown to translocate to the upper airways through sub-clinical micro-aspiration. Our findings reveal that *Pg*, upon translocating to this new niche, significantly attenuated lung damage by reducing viral loads during respiratory viral infections in the lungs of wild-type mice. This protective effect was attributed to the activity of gingipains, cysteine endopeptidases produced by *Pg*, which cleaved envelope glycoproteins on RSV as well as on related murine-specific Sendai virus (SeV), thereby impairing their infectious capacity. Notably, the reduction in viral loads was independent of interferon lambda (IFN-λ) signaling, which is actively suppressed by *Pg* in airway epithelial cells. However, the complete absence of IFN-λ signaling resulted in a stronger inflammatory response despite a low viral load. Thus, we show a previously undescribed role for the oro-respiratory bacterial colonizer *Pg* in creating bottlenecks to viral infection by the activity of its proteases.

**SIGNIFICANCE STATEMENT:** Reciprocal interactions between microbial colonizers and host epithelial cells are critical for providing initial defense against viral infections. However, our understanding of this phenomenon has been limited to microbiota-derived ligands that activate host pattern recognition receptors (PRRs), inducing basal interferon expression and downstream antiviral genes. Here, we present a novel mechanism that relies on microbial proteases to directly reduce viral load. Specifically, we discovered that the infectious capacity of the Respiratory Syncytial Virus (RSV) was significantly inhibited upon contact with the proteases (gingipains) produced by the oropharyngeal colonizer *Porphyromonas gingivalis*. Gingipains caused proteolytic degradation of the RSV envelope and attachment proteins, rendering them inactive. This preemptive reduction in viral infectious capacity consequently diminished the severity of respiratory viral infections in an IFN-independent manner.

## INTRODUCTION

Respiratory syncytial virus (RSV), a member of the *Pneumoviridae* family, is a significant human pathogen associated with considerable morbidity and mortality in infants and the elderly. It is the second leading cause of lower respiratory tract infections in children, with about 3% of those under the age of two requiring hospitalization (1), and also poses a significant health threat, particularly in the elderly, who also have a higher prevalence of periodontitis (2, 3). Despite the high incidence of infection and the development of RSV vaccines, immunity seems to wane over the lifetime, and RSV infections still bear a high risk for susceptible individuals (4). Thus, understanding the risk factors that modulate host susceptibility to RSV infections is essential for preventing severe disease.

The host microbiome has been shown to play a significant role in modulating host susceptibility to viral mechanisms, although the underlying mechanisms are complex, resulting in varied outcomes. For instance, the host microbiota is crucial for developing, priming, and maintaining innate and adaptive antiviral immune responses that protect against systemic viral infections. This protection is primarily provided by the detection of microbial metabolites or ligands from commensals by host pattern recognition receptors and downstream signaling that induce tonic levels of interferons and ISGs, which preemptively block viral infections (5–7). Another level of regulation arises from direct physical interactions between invading viruses and barrier microbiota. These interactions can promote viral infection by promoting viral uptake or cause viral inactivation by destabilizing viral particles. Thus resulting in synergistic or antagonistic outcomes (7). A growing body of evidence shows that in humans, the upper airways are densely colonized by bacteria very early in life. Microbiome profiling of the upper airways, throat, and nasopharyngeal surfaces in humans showed a dominance of oral biofilm bacteria within these passages (8–10). The nature of interactions between upper airway bacterial colonizers and invading respiratory viruses and their impact on respiratory viral infections are incompletely understood.

The oral cavity contains a rich and diverse microbiome that includes commensals as well as opportunistic pathogens capable of modulating host immune responses (11). Dysbiotic shifts in the oral cavity are associated with periodontitis, a destructive chronic inflammatory disease that has recently been linked with increasing the severity of multiple lung diseases such as emphysema, chronic obstructive pulmonary disease (COPD) and also respiratory viral infections such as SARS-CoV-2 and influenza in susceptible individuals (12, 13). Due to its anatomical location, the oral cavity is a portal of entry for respiratory viruses and oral bacteria that can translocate to the oro-respiratory and oro-pharyngeal mucosal surfaces via swallowing, coughing, or micro-aspiration (14–16). Several studies now show that even in healthy individuals, the lung microbiome is dominated by immigrating oral bacteria (*Prevotella sp; Fusobacterium sp; Streptococcus sp; Corneybacterium sp; Porphyromonas sp;*) that colonize the anatomically contiguous upper and lower airways via subclinical micro-aspiration (17–19). The presence of *Porphyromonas sp* in human airways (18, 20, 21) drove our interest, given its role in modulating epithelial antiviral immunity (22).

*Porphyromonas gingivalis (Pg)* is a significant periodontal pathogen that typically resides in the highly anaerobic subgingival niche, and its virulence potential can be modulated by other closely residing bacterial partners within the subgingival biofilm (11, 23). *Pg* is particularly adept at immune subversion, and its virulence factors, such as *Pg* proteases, are associated with broad immune dysregulation in humans (24, 25). We recently showed that *Pg* is particularly adept at inhibiting epithelial interferon (IFN) responses and modulating host susceptibility to infection (22). Here, we interrogated the impact of *Pg* infection in modulating host susceptibility to respiratory viruses, including RSV. Concordant with our observations in the gingival epithelium, *Pg* infection led to a broad suppression of IFNs and ISGs but paradoxically protected against viral infection. This protection was imparted by *Pg* proteases called gingipains that rapidly inactivated the infectious capacity of RSV and closely related Sendai virus (SeV) particles, ameliorating the severity of infection in vivo. Thus, our observations show that *Pg* can protect against RSV infection in an interferon-depleted environment.

## RESULTS

### *P. gingivalis* suppresses IFN responses in airway epithelial cells

First, we determined the transcriptional responses of respiratory epithelial cells to *Pg* infection, including the impact on antiviral pathways. We used the A549 cell line, previously used to interrogate transcriptional responses elicited by bacterial pathogens commonly associated with respiratory infections (26, 27). Both naïve and IFN-λ primed A549 cells were infected with *Pg*, and transcriptional responses were determined by RNA-seq. We found that while *Pg* could attach and invade A549 cells (**Fig S1A-C**), *Pg* infection led to modest changes in gene expression (**Fig 1A**). *Pg* upregulated multiple genes involved in extracellular matrix remodeling and lung fibrosis (**Fig S1D-E**) but had a minimal impact on inflammatory and IFN-regulated pathways typically essential for orchestrating epithelial antimicrobial responses and immune cell recruitment in *vivo* (**Fig 1 A-B**). The transient colonization of mouse lungs with a low single dose of *Pg* (10^7^ CFU) intratracheally also had a minimal impact on body weight (**Fig S2 A-B**), recruitment of immune cells (**Fig S2 C**), and inflammatory cytokine gene expression (**Fig S2 D**). in the lungs of infected mice. This was unsurprising as *Pg* can be found in low numbers in the oral cavities of healthy individuals with no apparent signs of periodontitis (28, 29). It has also been found in the tracheal aspirates of healthy individuals (18, 20, 21), indicating that *Pg* colonization of the lung is not associated with adverse inflammatory impacts or disease. Next, we determined whether *Pg* colonization interfered with airway epithelial responses to a secondary challenge, such as IFN induction and priming that occurs during viral infection.

**Figure 1:**
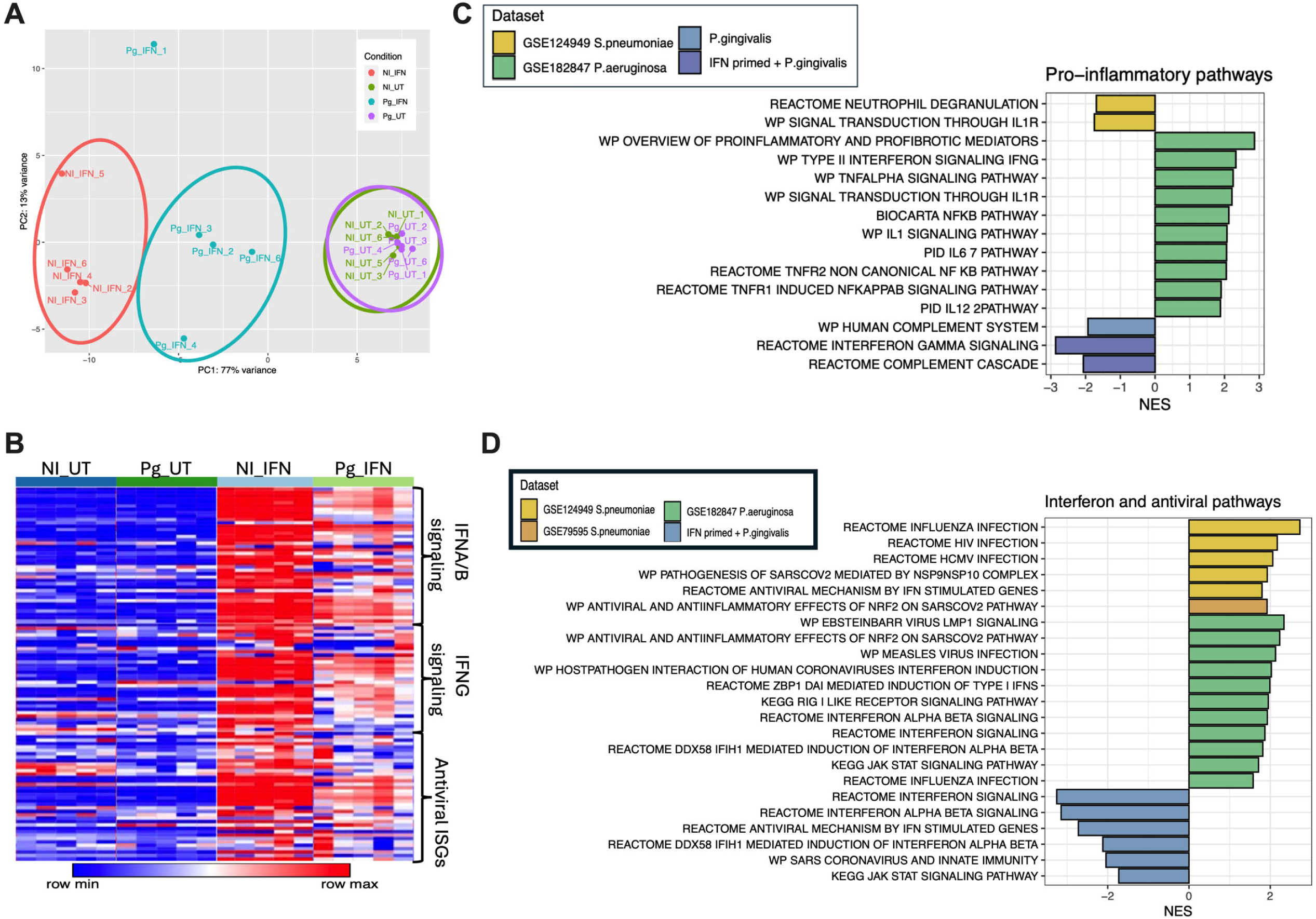
*Pg* infection does not elicit IFN responses in AEC: (A-B) Naive and IFN-λ primed (20 ng/ml; overnight) A459 cells were infected with *Pg* (MOI 100) for 24h, and transcriptional responses were analyzed by RNA seq. (A) Individual samples were categorized into four groups: untreated cells (NI_UT), IFN-primed cells (NI_IFN), *P. gingivalis*-infected cells (Pg_UT), and IFN-λ primed *Pg* infected cells (Pg_IFN). (B) Heatmap, showing log-normalized expression of genes from selected pathways: REACTOME_INTERFERON_ALPHA_BETA_SIGNALING (Stable Identifier: R-HSA-909733), REACTOME_INTERFERON_GAMMA_SIGNALING (Stable Identifier: R-HSA-877300), REACTOME_ANTIVIRAL_MECHANISM_BY_IFN_STIMULATED_GENES (Stable Identifier: R-HSA-1169410). (C) Pro-inflammatory or (D) IFN/antiviral pathways were generated from publicly available bulk RNA-seq datasets: A549 cell line infected with *S. pneumoniae* (GSE79595) or *P. aeruginosa* (GSE182847); nasal mucosal cells infected with *S. pneumoniae* (GSE124949).

We previously showed that *Pg* can actively suppress inducible IFN responses in gingival epithelial cells upon secondary challenge with an array of viral ligands and activated diverse pattern recognition receptors (PRRs) to induce IFNs (22). Thus, we determined whether *Pg* inactivated or interfered with the efficacy of IFN priming and ISG expression in airway A549 cells. Interestingly, we saw that *Pg* infection markedly suppressed IFN signatures in IFN-λ primed A549 cells, indicating that *Pg* upon invasion into epithelial cells can override the stimulative effects of IFN priming (**Fig 1A-B; Fig S1F-G**). We also observed refractoriness in IFN responses upon secondary challenge with Poly I: C (a viral mimetic) in A549 cells and other airway epithelial cell lines (**Fig. S1H**). This indicated that the IFN suppressive capacity of *Pg* was not limited to a single cell line. We compared the transcriptional signatures induced during *Pg* infection to publicly available RNA-seq datasets from other pneumonia-causing bacterial pathogens, *Streptococcus pneumoniae* and *Pseudomonas aeruginosa*. All datasets utilized the A549 cell line, except the *S. pneumoniae* dataset, which was from nasal epithelium harvested from healthy adult volunteers. Our data show that unlike *Pg*, *S. pneumoniae and P. aeruginosa* upregulated inflammatory signaling pathways and/or IFN signaling in airway epithelial cells (**Fig 1C, D**). Thus, *Pg* was unique in its capacity to strongly suppress IFNs in the airway epithelium, possibly predisposing it to viral infection.

### *P. gingivalis* co-infection limits IFN responses to RSV

To determine how *Pg* impacted RSV infection, we used a model of air-liquid interface cultures containing human bronchial epithelial cells (HBE) generated by the expansion of human bronchial basal cells in transwells (30) (**Fig. 2A**). Well-differentiated HBE cultures contain ciliated epithelial cells, basal cells, and goblet cells that naturally produce mucus (see H&E and PAS staining (**Fig. S2 E-F**) and thus represent the physiological microenvironment of the human lung. HBE transwells were infected with the rgRSV224 strain (derived from RSV A2 strain), containing an inserted green fluorescence protein (GFP) gene before the first gene in the full-length cDNA copy of the RSV genome (31). GFP fluorescence thus serves as a ‘tracer’ for quantitatively measuring RSV infection and replication. RSV infection of HBE cultures strongly induced high levels of IFN-λ (**Fig. 2B**) and, to a lower extent, IFN-β (**Fig. 2C**); however, both were significantly suppressed in the presence of *Pg*. We were unable to detect IFN-α or IFN-γ by ELISA. *Pg* produces three different endoproteases with trypsin-like activity called gingipains. Arginine gingipains (HRgpA and RgpB) and lysine gingipains (Kgp) cleave after the (Arg-X-aa) and Lys-X-aa peptide bonds, respectively (32, 33). We previously showed that all 3 gingipains drive the proteolysis of the type I and III IFN receptors, blocking downstream ISG activation in the oral epithelium (22). RSV coinfection with WT, Δ*rgpa,* Δ*rgpb,* Δ*kgp* strain (abbreviated as Δ*KRAB*) (34) and the invasion deficient Δ*fimA* mutant (35) showed that gingipains activity was essential in limiting the efficacy of IFN signaling and downregulating ISG expression (**Fig S2 G**). Interestingly, despite the trypsin-like activity of gingipains, we did not observe any cellular sloughing or breakdown of HBE layers in *Pg* co-infected wells (**Fig 2D**). Vacuolation and stress responses in HBE layers, typical of RSV infection, were more evident in RSV infected wells, indicating that *Pg* counterintuitively protected against RSV-mediated pathology by a mechanism distinct from IFN suppression.

**Figure 2:**
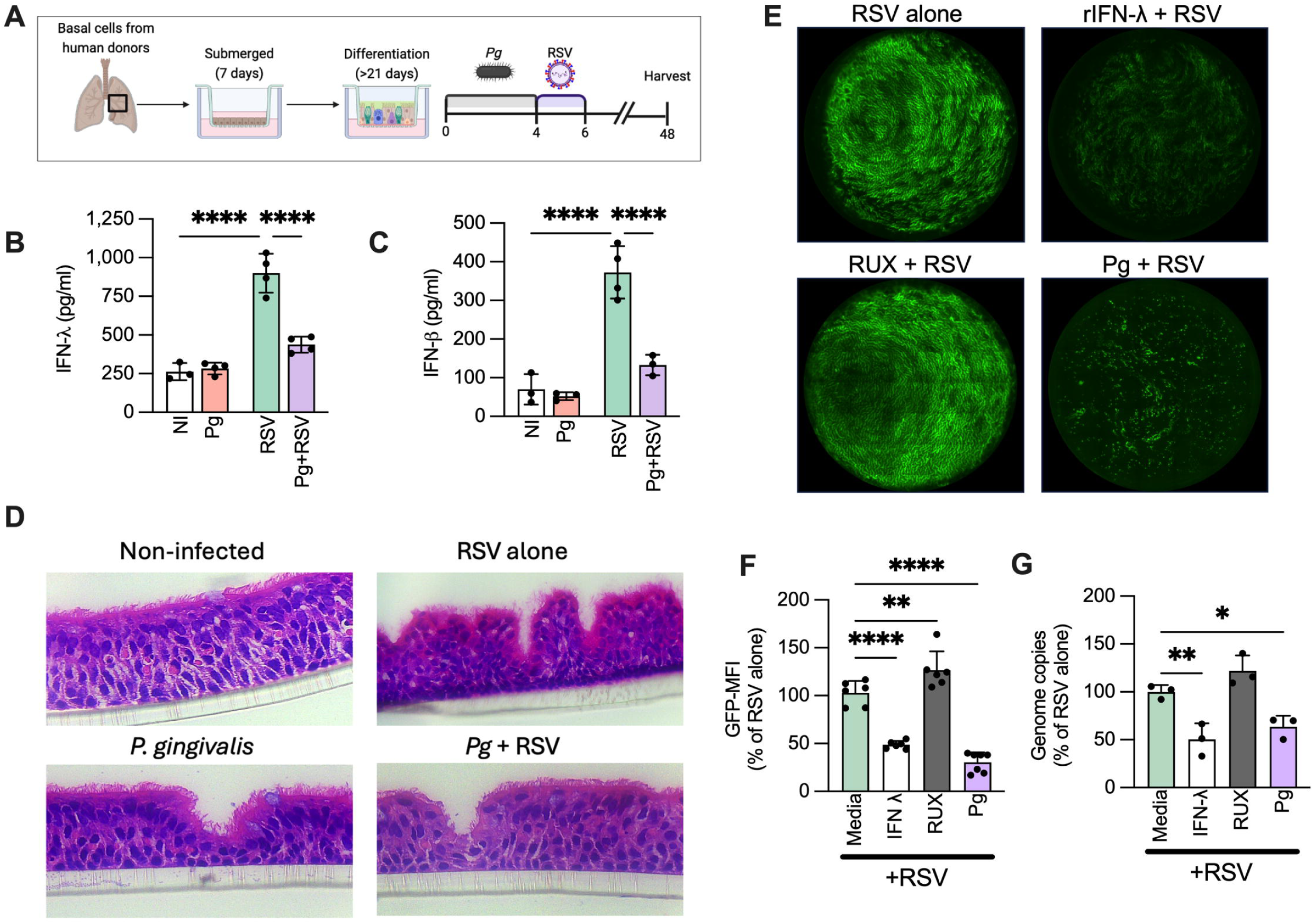
*Pg* restricts IFN-λ responses to RSV infection: (A) Graphical depiction of the air-liquid interface human bronchial epithelial (HBE) transwell cultures and differentiation and infection. Transwells were infected with *Pg* (5x10^6^CFU) for 4 h and then by rgRSV224-GFP strain (∼1300 PFU) for 2 h, and incubated at 37°C for an additional 48 h. (B) IFN-λ and (C) IFN-β levels in basal media 48 h post-RSV infection (mean ± SD). Statistical differences were calculated using Two-way ANOVA (****P<0.0001). (D) Representative H&E images of HBE membranes at 48 h. HBE transwells were incubated with either 20 ng/ml IFN-λ for 24 h or 10µM ruxolitinib (RUX) prior to RSV infection. (E) Representative images of RSV-infected HBE transwells showing fluorescent infected cells 48 h post-infection. (F) Average GFP-MFI from 3-6 wells per group is shown as mean ± SD. (G) Average genome copies of RSV were enumerated by qPCR 48h post-infection and shown as mean ± SD. Statistical differences for F-G were determined by one-way ANOVA (*P<0.05; **P<0.01; ****P<0.0001).

### Despite the absence of IFNs, RSV infection is counterintuitively suppressed by Pg

RSV is an IFN-restricted virus, and several IFN-induced antiviral restriction factors, such as the IFN-induced transmembrane proteins (IFITM) 1 and 3, inhibit RSV entry and replication. Their deletion in mice was associated with significant disease and weight loss (36, 37). Other IFN-induced proteins (IFI) 44 and 44L have also been shown to restrict RSV replication and transcription (38). Consistent with this, we observed that the pre-treatment of HBE transwells with rIFN-λ significantly blocked RSV replication, and pre-treatment with ruxolitinib (RUX), a JAK1/2 inhibitor that blocks IFN-activated ISG expression, augmented RSV replication (**Fig. 2E-F**). Since *Pg* infection creates an IFN-depleted microenvironment (**Fig 1, 2**), we expected a significant increase in RSV replication in *Pg* co-infected cells. However, contrary to our expectation, RSV replication (GFP-MFI and genome copy numbers) was significantly dampened in *Pg* co-infected wells (**Fig. 2 F-G**).

### Gingipains strip viral glycoproteins, reducing their infectious capacity

Next, we explored whether *Pg* virulence factors impacted RSV infectivity or replication during co-infection. Respiratory viruses are particularly adept at co-opting host proteases for entry into host cells, virion maturation, and modulating host response during infection (39). However, the impact of proteases emanating from bacteria colonizing the on regulating virion infectivity has not been described. In the HBE model, focus-forming units (FFUs) correlate with the number of infection foci created upon virion binding or the total number of infected cells based on fluorescently labeled viral antigens for non-lytic viruses. In the RSV model, GFP FFUs at 24 h indicate the number of infectious viral particles. At later stages (48 h post-infection), GFP mean fluorescence intensity (MFI) increases due to viral replication and syncytia formation due to ciliary beat characteristic of HBE cultures (40). We observed a significant drop in RSV infectivity (FFUs) in cells co-infected with *Pg* WT at 24h post-infection but not with the Δ*KRAB* mutant, indicating that *Pg* inhibited adsorption of the virus in a gingipain-dependent manner (**Fig 3A, B**). This initial loss of infection load correlated with reduced GFP MFI at 48 h (**Fig. 3C, D**). The invasion deficient Δ*fimA* mutant, which lacks the major fimbrial antigen FimA but has intact gingipain activity, showed a similar phenotype at the wildtype strain of *Pg*.

**Figure 3:**
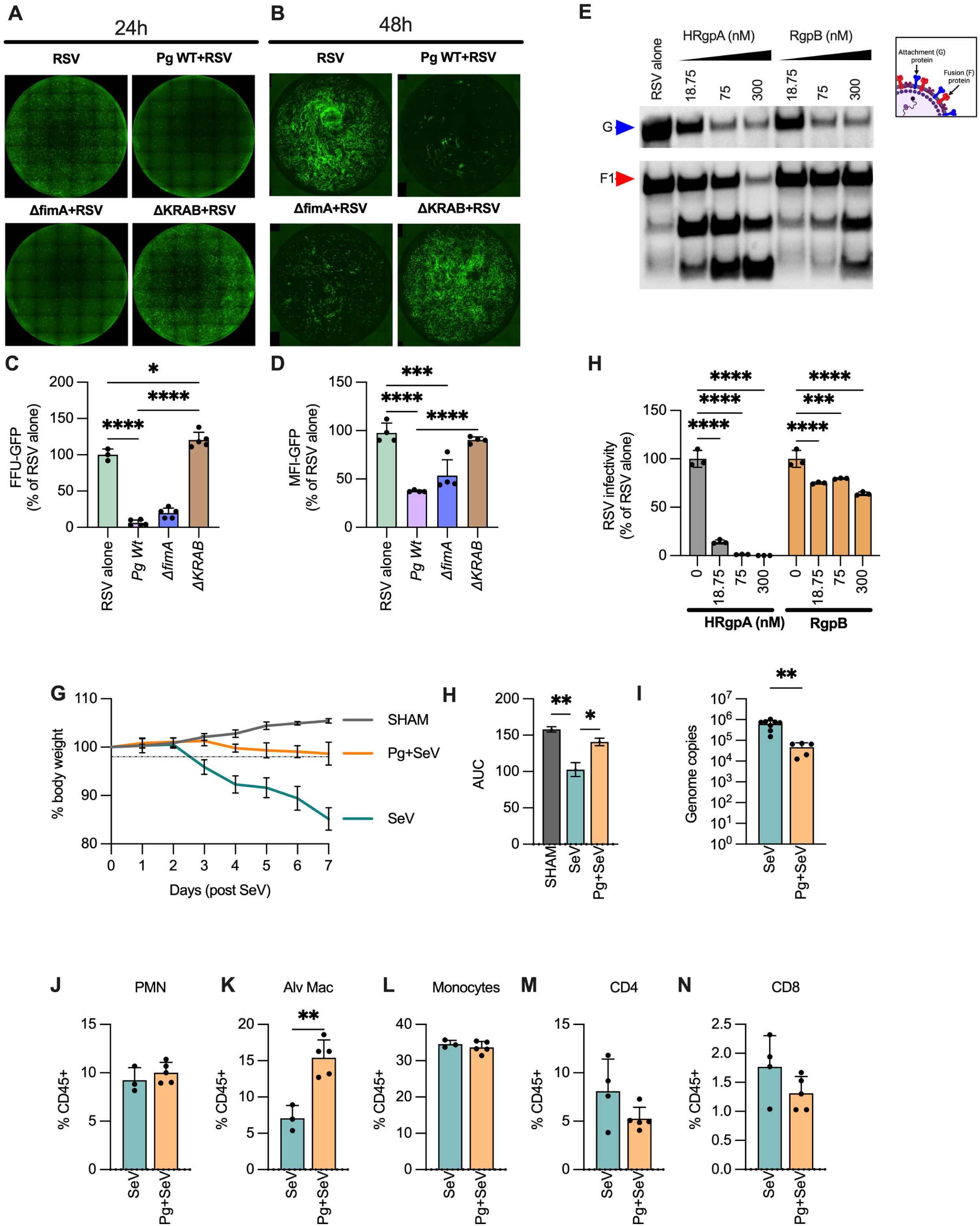
Gingipains impair RSV infectivity: (A-D) HBE cultures were infected with 5x10^6^ CFU of wildtype (WT) *Pg* or isogenic mutants Δ*fimA* and Δ*rgpA,* Δ*rgpB,* Δ*kgp* triple mutant (abbreviated as Δ*KRAB*) for 4 h, followed by a 2 h infection with RSV (∼1300 PFU). Representative images of RSV infection at 24h and 48 h post-infection. Average mean fluorescence intensity (MFI) from 3-4 wells per group is shown as mean ± SD. Statistical differences were determined using one-way ANOVA (*P<0.05; ***P<0.0005; ****P<0.0001). (E) 1x10^6^ RSV particles were treated with arginine gingipains HRgpA and RgpB at indicated concentrations for 30 mins, and RSV attachment (G; blue arrow) and fusion (F; red arrow) protein integrity (laddering) were determined by a western blot. (H) Hep2 cells were infected with gingipain-treated RSV particles, and infection was determined by quantifying GFP fluorescence 24 h post-infection. Data was normalized to RSV alone (no gingipain treatment) and is shown as mean ± SD. Statistical differences were determined using two-way ANOVA (****P<0.0001). (G-N) WT mice were colonized with 10^7^ CFU *Pg* intratracheally and 24 h later challenged intranasally with 2x10^5^ PFU Sendai virus (SeV). (G) Weight loss over 7 days is shown as % of initial body weight and (H) area under the curve (AUC) as mean ± SD. Statistical differences were measured using one-way ANOVA (*P<0.05, **P<0.01). (I) Viral genome copies (mean ± SD) in the lung homogenates of mice on Day 7. (I-N) Immune cell infiltration in mouse lungs (4-5 mice) was determined by flow cytometry, and population abundance is shown as % CD45 (mean ± SD). Statistical differences were calculated using a t-test (**P<0.005).

RSV encodes a glycoprotein (G) known as an attachment protein and a fusion (F) protein that play important roles in host cell entry. Unlike cell lines, where RSV entry relies on G protein binding to cell surface glycosaminoglycans (GAGs), and in particular heparan sulfates, infection in HBE cultures occurs exclusively by binding to ciliated cells in a CX3CR1-dependent manner (41, 42). Since eliminating gingipains restored RSV infectivity in *Pg* co-infected cells, we determined whether gingipains incapacitated virions by cleaving RSV envelope glycoproteins. Preincubation of RSV with purified Arg gingipains led to the proteolysis of RSV attachment (G) and fusion (F) proteins (**Fig. 3E**). Since G and F proteins are essential for viral entry into the host cells, we pre-treated RSV with Arg gingipains and determined their infectious capacity on HEp2 monolayers. We saw a significant drop in RSV infectivity (**Fig. 3F**), confirming that gingipain mediated cleavage of F and G proteins dramatically lowered the infectious capacity of RSV virions.

### *Pg* colonization of upper airways attenuates viral loads *in vivo*

RSV is a human-adapted virus that poorly infects mice (C57BL6/J strain) at physiological doses (43, 44). Thus, we used Sendai virus (SeV), a negative-sense single-stranded RNA virus belonging to the paramyxoviridae family of viruses and specifically infects rodents. SeV infection in mice parallels the clinical symptomology associated with RSV infection in humans (45). Interestingly, similar to our observations with RSV, gingipain treatment caused the proteolysis of the hemagglutinin-neuraminidase (HN) protein and fusion (F) protein essential for SeV infection (**Fig. S3 A**).

To determine the impact of *Pg* colonization of SeV infection in *vivo*, we first colonized mice with *Pg* intratracheally and then, 24 hours later, challenged them intranasally with SeV. While SeV infection led to significant weight loss, *Pg*-SeV co-infected mice showed a marked attenuation in weight loss, which correlated with a significant reduction in viral genome copies in the lungs of SeV-infected mice (**Fig. 3G-I**). SeV infected mouse lungs showed characteristic diffuse damage to the alveolar epithelium and immune cell infiltration. In contrast, *Pg-SeV* co-infected mice seemed to attenuate signs of damage and inflammation (**Fig. S3 B**). However, analysis of lung single-cell suspensions showed no statistical differences in inflammatory cell infiltration (**Fig. 4J-N**; see gating strategy in **Fig. S3 C**) in SeV alone and *Pg* + SeV co-infected mice, indicating that the protection conferred upon introduction of *Pg* in mouse airways was unrelated to differential immune cell recruitment but more likely related to the physical interaction of *Pg* with the virus and attenuation of viral load.

**Figure 4:**
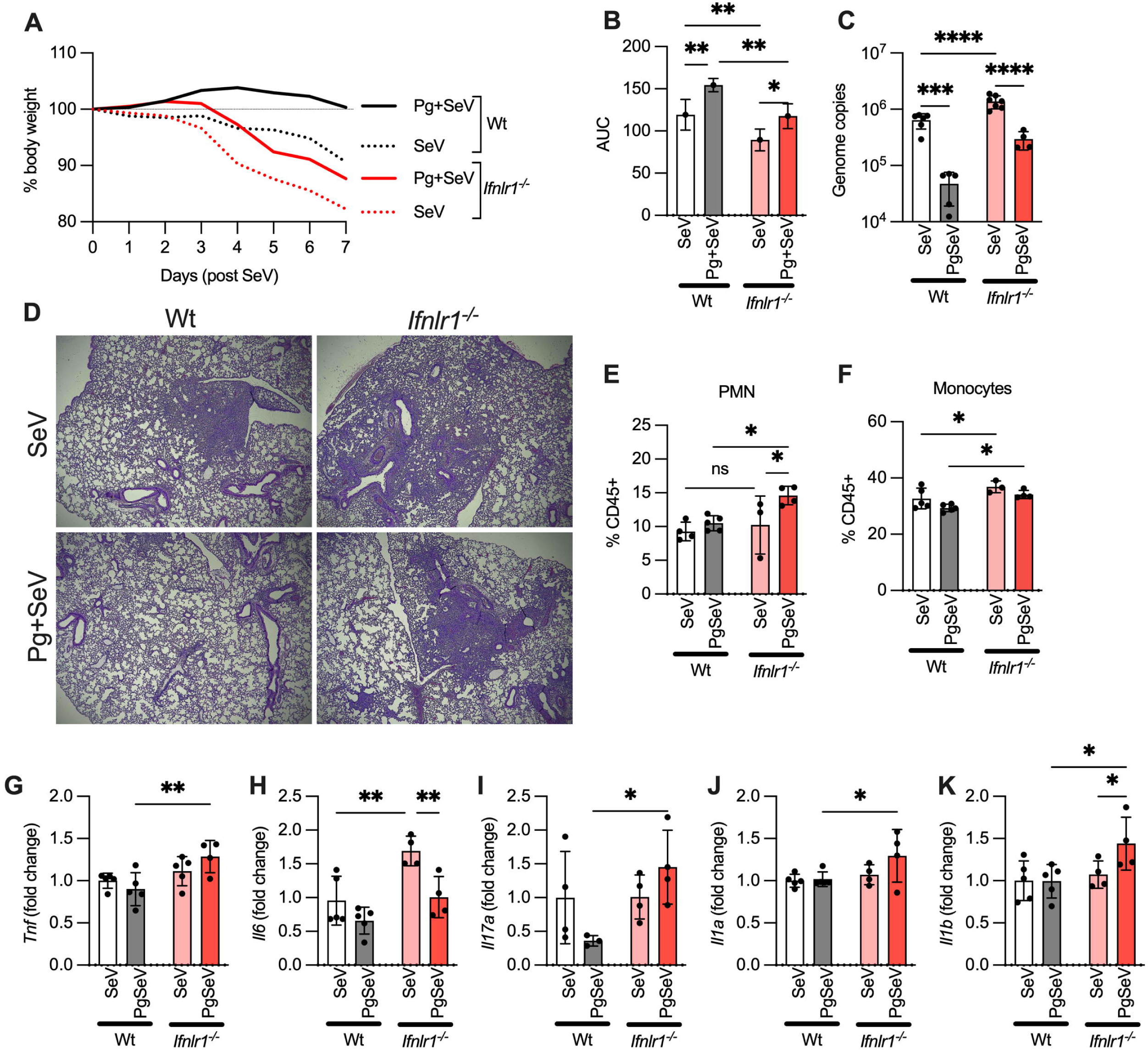
*Pg*, infection in *Ifnlr^-/-^* mice is associated with significant lung damage: WT and *Ifnlr^-/-^* mice were colonized with 10^7^ CFU *Pg* intratracheally and 24 h later challenged intranasally with 2x10^5^ PFU Sendai virus (SeV). (A) Weight loss over 7 days is shown as % of initial body weight and (B) area under the curve (AUC) as mean ± SD. (C) Viral genome copies (mean ± SD) in the lung homogenates of mice on Day 7. (D) Representative images of H&E-stained lung tissues on Day 7. (E-F) Flow staining showing relative % of neutrophils (PMNs) and monocytes in mouse lungs on Day 7. (G-K) inflammatory cytokine transcripts were assessed by qPCR and normalized to GADPH. For G-K, data are shown as fold change over the WT-SeV group. Statistical differences were measured using Two-way ANOVA (*P<0.05, **P<0.01; ****P<0.001).

### *Pg* co-infection in *Ifnlr^-/-^* mice exacerbated SeV induced lung damage despite a reduction in viral loads

*Pg* is a human-adapted pathogen that can be acquired within the first few days of life in humans and is also detected in periodontally healthy individuals, albeit at a lower frequency compared to patients with periodontitis (28, 29). We previously showed Pg abundance in the gingival tissues of patients with periodontitis correlated with suppression in IFN responsiveness (22). However, mice are not a natural host for *Pg;* unlike humans, mouse airways are unlikely to be continuously seeded naturally via aspiration from the oral cavity. In our mouse model, intratracheal delivery of Pg into mouse airways efficiently attenuated the infectivity of SeV in vivo; however, it did not reproducibly block IFN expression. To overcome this limitation and determine the combined impact of *Pg* colonization and IFN suppression in modulating host responses to respiratory viruses, we used IFN-λ receptor knockout mice (*Ifnlr^-/-^*). Historically, IFN-λ has been shown to be essential in restricting initial viral infection and spread without causing damaging inflammation or immunopathology typically associated with Type I IFN signaling (46, 47).

We first intratracheally colonized WT and *Ifnlr ^-/-^* mice with *Pg*, followed by intranasal SeV challenge after 24 h. *Ifnlr ^-/-^* mice are more susceptible to SeV infection and have significantly higher weight loss over 7 days with higher viral titers in lung homogenates than wildtype mice (**Fig 4 A-C**). *Pg* could only modestly attenuate viral loads and weight loss in *Ifnlr ^-/-^* mice, indicating that *Pg* infection of the airways creates a bottleneck for viral infection, reducing SeV titers independent of IFN-λ signaling (**Fig 4C**). However, despite the attenuation in viral loads, *Pg*+ SeV co-infected *Ifnlr ^-/-^* mouse lungs showed significant damage with inflammatory cell (neutrophils and monocyte) recruitment (**Fig 4 D-F**) compared to WT mice. Previous studies have shown that IFN-λ signaling during viral infection was essential in regulating neutrophil effector functions and damage (48). Consistent with this, we saw significantly higher infiltration of neutrophils and monocytes into the lungs of *Ifnlr ^-/-^* mice, which correlated with the pathological damage in the lungs of SeV-infected mice (**Fig. 4D-F**). Interestingly, the introduction of Pg into the airways almost entirely blocked inflammatory damage in the lungs of wildtype mice but not *Ifnlr ^-/-^* mice (**Fig. 4D**) despite the reduction in viral load (**Fig. 4C**). qPCR analysis of lung homogenates also showed significantly higher levels of multiple pro-inflammatory cytokine transcripts (**Fig 4G-K**) consistent with immune cell infiltration and damage. Overall, these data shows that the presence of *Pg* in the lungs of immunocompromised mice can attenuate viral loads but also contribute towards more damage in respiratory viral infection.

## DISCUSSION

Here, we demonstrate the direct role of bacterial proteases (gingipains) in decreasing the infectious capacity of respiratory viruses, thereby modulating host susceptibility to infection. This mechanism of viral-bacterial antagonism is distinct from previously described roles for the host microbiota in regulating host susceptibility to viral infection and identifies a new role for oro-respiratory microbial colonizers in limiting the infectious dose of a respiratory virus in a manner that is independent of IFN signaling.

Independent of host antiviral priming (5), recent studies show that physical interactions between viruses and bacteria can also significantly impact host susceptibility to viral infection by directly affecting virion stability, cellular entry, and the multiplicity of infection, ultimately impacting the infectious dose of invading viruses. For example, the binding of capsid-1 protein from the enteric poliovirus to bacterial cell wall components such as lipopolysaccharides (LPS) or peptidoglycans (PG) was shown to enhance the stability of the poliovirus particles, resulting in a higher infection burden and transmissibility (49). Viral adsorption on bacterial adhesins can also enhance the multiplicity of infection by phagocytic uptake of viral-bacterial complexes. Thus, in the cases mentioned above, the gut microbiome bacteria play more of a ‘pro-viral’ role in enhancing host susceptibility to viral infection (50, 51). In contrast, antagonistic interactions can also occur. For example, *Lactobacillus reuteri,* a commensal, can inhibit viral entry into host cells by binding and sequestering the virus away from the host cells (52). We showed that purified gingipains rapidly inactivate gingipains even in the absence of live *Pg*. Thus, our findings are another layer to this complexity by demonstrating that secreted proteases by a bacterium can also modulate viral fitness even without direct bacterial-viral interactions.

Arginine and Lysine gingipains together account for over 80% of the proteolytic activity of *Pg,* and their trypsin-like activity has been associated with the proteolytic degradation of numerous cellular proteins, receptors, cytokines, adhesion molecules, immunoglobulins, proteins in the complement and coagulation cascades, and extracellular matrix proteins (32, 33). Drugs specifically designed to target cationic arginine residues on viruses such as SARS-CoV-2, RSV, and influenza (H1N1) have been modeled to interfere with viral entry into cells, highlighting the significance of arginine-based interactions in viral infection (53). Although we primarily focused on RSV and SeV in this paper, given the ability of gingipains to target arginine and lysine residues, it may protect against a broad range of viruses, possibly via the degradation of other viral envelope or spike proteins. Gingipains are secreted by *Pg* and can be found not only in the oral cavity but also in other organs (54). Studies in periodontitis patients show that their concentration in the gingival crevicular fluids of periodontitis patients may reach as high as ∼100 nM (55). Thus, they may still be able to inactivate viruses in these patients despite the dampened IFN signaling induced by *Pg*.

Finally, our data shows that gingipain activity was protective during viral infection despite the significant loss of IFN-λ. HBE cultures and lungs of wildtype mice co-infected with *Pg* showed significantly reduced damage, concomitant with viral clearance and/or reduced infection. Recent studies show that IFN-λ signaling might play a more nuanced or context-dependent role in tissue repair following viral infection. Two recent reports show that IFN-λ signaling in the lungs during high-dose IAV infection or repeated Poly IC instillation counterintuitively hampered lung repair by inhibiting epithelial proliferation and differentiation (56, 57). Lowered expression of IFN-λ during *Pg* infection might lead to a fine-tuned or regulated response that protects barrier integrity during RSV/ SeV co-infection, resulting in reduced damage. Interestingly, low-dose instillation of *Pg* in mouse lungs did not induce inflammation and was efficacious in reducing viral titers in vivo and protecting against lung damage. Thus, we show a novel role for an oral bacterium and in protection against respiratory viral infections and tissue damage.

## MATERIALS & METHODS

### Bacteria

*P. gingivalis* 33277 (wild type) was cultured in trypticase soy broth (TSB) supplemented with 5 μg/mL hemin, 1 μg/mL menadione, and 1 mg/mL yeast extract. Isogenic mutants were cultured with selection antibiotics: 10µg/ml erythromycin for major fimbriae mutant (Δ*fimA*)(58); 10 μg/mL erythromycin, 0.7 μg/mL tetracycline, and 20 μg/mL chloramphenicol for triple gingipain mutant Δ*kgp* Δ*rgpA* Δ*rgpB* abbreviated as: (Δ*KRAB*)(34, 59). HaloTag *Pg* (ΔInlJc/TCOW-Halo-InlJ) was made by conjugation of the Δ*InlJ* mutant with the S-17 strain of *E. coli* harboring the pT-COW plasmid containing Halo-InlJ. HaloTag-*Pg* was cultured with 10µg/ml erythromycin and 0.7 μg/mL tetracycline. All strains were grown anaerobically (85% N_2_, 10% H_2_, and 5% CO_2_) at 37 °C.

### Virus production and quantification

Recombinant GFP-expressing rgRSV224 is based on the A2 laboratory strain of RSV that carries a GFP tag on the leader sequence of the virus (60). RSV stocks were grown in HEp-2 cells in EMEM-10% FBS. GFP-expressing viruses were serially diluted and titrated on HEp-2 cells by inoculating for 2 h at 37°C and counting GFP foci at 24 hp. RSV genome copies in infected HBE cultures were determined by qPCR using the Power SYBR Green PCR Master Mix (ThermoFIsher) with primers designed to target the nucleocapsid gene. Forward primer: 5’-GGGAGAGGTAGCTCCAGAATA-3’, and reverse primer sequence: 5’-CTCCTAA TCACGGCTGTAAGAC-3’. Copy numbers were determined using a standard curve of Quantitative Genomic RNA from human RSV strain A2 purchased from ATCC. Sendai virus (strain 52) was purchased from ATCC (cat. number: VR-105). SeV genome copies in mice were quantified using SeV standard as previously described (61). SeV qPCR was performed using TaqMan mastermix and the N3F gene SeV-1237F (5’ GGCGGTGGTGCAATTGAG 3’), N3R gene SeV-1300R (5’ CATGAGCTTCTGTTTCTAGGTCGAT 3’) and MGB probe N gene SeV-1257 (5’ AGCTGTAGACAATGCC 3’) probe sets.

### SEV infection in mice

6-10-week-old C57BL/6 mice were purchased from Jackson Laboratories (Bar Harbor, ME). Intratracheal instillation of 10^7^ CFU of *Pg* was done using a catheter in intubated anesthetized mice. After 24h, mice were challenged with 2x10^5^ PFU of SeV intranasally. Body weights were measured daily for 7 days. Tissues were harvested on day 7 post-SeV infection.

### Primary human bronchial epithelial culture and differentiation (HBE) and infection

HBE progenitors were isolated from human donor airways as previously described (30). For setting up HBE transwells, progenitor cells were plated on 0.4 μM pore Corning transwells membranes (6.5 mm or 12 mm in diameter) and fed with PneumaCult air-liquid interface (ALI) media (Stemcell Technologies) supplemented with ROCK inhibitor in both the apical and basolateral chambers. Media in both chambers was replaced every 2–3 days. At 7 days, when the cells were confluent and had formed tight junctions as demonstrated by electrical resistance, the apical medium was removed, and the basal medium was replaced with Pneumacult-ALI Medium (STEMCELL Technologies). The media in the basal chambers was replaced with fresh media every 2-3 days, and the apical surface was washed with 100 μL of DMEM to remove accumulating mucus for 4 weeks, which is sufficient time for complete differentiation of HBE (40). For all experiments, mature HBE cultures were infected with *Pg* (5x10^6^ CFU) for 4h on their apical surface. After infection, the apical bacteria-containing fluid was removed and replaced with rgRSV224 inoculum (1300PFU) for 2 h at 37°C. At the end of this period, the apical fluid was removed, and cultures were incubated at 37°C. For specific experiments, HBE cultures were treated on the basal side with recombinant 20 ng/ml human IFN-λ1 (Peprotech) or 10 µM Ruxolitinib (Invivogen) before bacterial infection.

### Statistics

Statistical analyses utilized GraphPad Prism 6.0 (GraphPad). A *P* value <0.05 was considered statistically significant. A detailed description of the statistical tests used is stated in each figure legend.

*Additional methods are listed in the Supplemental DATA

## AUTHOR CONTRIBUTIONS

CJRH carried out all experiments, analyzed data, and wrote the manuscript with input from RJL and JB. MT and MNA analyzed all RNAseq datasets; BP and JP provided gingipains and Pg mutants and expertise on gingipain biology; PC, MEP and MHG provided reagents and expertise with viral models; JP and RJL provided *Pg* strains and KAC helped with mouse infections.

## ACKNOWLEDGEMENTS

These studies were supported by DE011111 to RJL and DE031756 to JB and RJL.

