## Supplemental Methods and Figure Legends for "Virion stripping: A new role for bacterial proteases"

**Supplemental Figure Legends:**

**Supplemental Figure S1:** (A-C) HEp2 cells were infected with Halo-tagged labeled *Pg* 33277. *Pg* infected cells or TMR dye-positive cells were identified by flow cytometry (‘Total *Pg*’ gate). Invasion vs. adsorption was determined based on the accessibility of anti-*Pg* antibody from the total *Pg* gate and quantified as described in methods. (D-E) Naïve A549 cells were either left untreated (NI_UT) or infected with *Pg* (MOI 100) for 24 h (Pg_UT), and transcriptional responses were analyzed by RNA seq. Heatmaps, showing log-normalized expression of genes involved in selected pathways: WP_LUNG_FIBROSIS (<https://www.wikipathways.org/pathways/WP3624.html>), REACTOME_EXTRACELLULAR_MATRIX_ORGANIZATION (Stable Identifier: R-HSA-1474244) Color intensity denotes the level of gene expression. (F-G) Volcano plots showing differential expression of genes. The x-axis represents log2 (fold change) of gene expression, and the y-axis represents log-transformed P-value. The red and blue dots indicate the significantly upregulated and downregulated genes, respectively. The grey dots indicate genes that did not meet the cut-off criteria for significance in differential expression. (H) A549, Calu3 or HEp2 cells were infected with *Pg* (MOI 100) for 4 h and then challenged with Poly I:C overnight. IFN-λ production is shown as mean ± SD. Statistical differences were measured using Two-way ANOVA (****P<0.0001).

**Supplemental Figure S4:** (A) To assess the impact of gingipains on SeV infectivity, 1x10^6^ SeV particles were treated with different concentrations of the arginine gingipains (HRgpA and RgpB), and lysine gingipain (Kgp) for indicated time points. Western blot image showing degradation (laddering) of SeV hemagglutinin neuraminidase (HN) and fusion (F) protein. (B) Representative H&E staining of lungs from WT mice colonized with 10^7^ *Pg* intratracheally and subsequently (24 h later) intranasally with 2x10^5^ TCID_50_ of SeV 7 days post-infection. (C, D) Gating strategy for the analysis of leukocytes (CD45^+^) in single cell suspension of lungs 7 days post-infection. B cells (CD45^+^, CD19^+^); alveolar macrophages (CD45^+^, CD11c^+^, SiglecF^+^); eosinophils (CD45^+^, Siglec F^+^); monocytes (CD45^+^, Ly6C^hi^); neutrophils (Ly6G^hi^, Ly6C^int^); CD4 T cells (CD45^+^, TCRb^+^, CD4^+^) and CD8 T cells (CD45^+^, TCRb^+^, CD8^+^).

**SUPPLEMENTAL METHODS:**

*Mouse tissue processing:* Mouse lung lobes were harvested and snap frozen for RNA isolation, fixed in 10% NBF for histology, or enzymatically digested to obtain a single cell suspension for flow cytometry. Briefly, the lung lobes were cut into small pieces and digested using 250 U/ml collagenase IV, 5 U/ml hyaluronidase, and 50 U/ml DNase at 37°C. After 45 min, 10mM EDTA (final concentration) was added, and tissues were incubated for an additional 15 min at 37°C to stop enzyme activity. Cell suspensions were strained using a 70µm strainer and stained for flow cytometry using anti-mouse CD45-BUV737, CD19-FITC, CD11c-Alexa fluor 647, SiglecF-PE, Ly6C-BV785, Ly6G-BV650, TCRβ-FITC, CD4-BV510 and CD8-APC-H7 on ice for 45 min. Cells were washed in flow buffer (PBS, 2 mM EDTA, and 0.5% BSA). Samples were run on an LSR Fortessa X-20 (BD), and data was analyzed using Flowjo software.

*Cell Culture and Infection:* The human alveolar basal epithelial A549 cells were cultured in Kaighn’s Modification on Ham’s F12 medium supplemented with 10% heat-inactivated fetal bovine serum (ΔFBS). Calu-3 lung adenocarcinoma cell line was cultured in Eagle’s Minimum Essential Medium (EMEM) supplemented with 10% ΔFBS. HEp-2 cell line was cultured in Dulbecco’s Modified Eagle’s Medium (DMEM) supplemented with 10% ΔFBS. All cell lines were grown in a 37 °C incubator and 5% CO_2_. 70-80% confluent monolayers were challenged with *Pg* (mid-log phase) at a multiplicity of infection (MOI) of 100 bacteria per cell for 4 hours at 37°C. Cells were washed with PBS wash to remove uningested bacteria. In certain experiments, cells were challenged with Poly I:C/LyoVec low molecular weight stimulation (500ng/ml) for 24 hours after *Pg* infection.

*Pg attachment and invasion of airway epithelium:* Mid growth phase HaloTag *Pg* was stained with 6.25 nM HaloTag TMR ligand (Promega) for 1 h at room temperature, followed by 2 washes in PBS. A549 cells were infected with pre-stained HaloTag *Pg* at a MOI of 100 for 4 hours at 37°C. After infection, cells single cells were dislodged using Trypsin-EDTA (0.05%), Fc blocked, and stained with anti-*Pg* 33277 rabbit immune sera (1:500) (Sztukowska et al., 2016) for 45 min at 4°C. Cells were washed twice with flow buffer and then stained with Alexa Fluor 488–conjugated anti-rabbit secondary antibody at 1:1000 dilution for 1 h and analyzed by flow cytometry using an LSR Fortessa X-20 cytometer (BD).

*Gingipain treatment of respiratory viruses:* Gingipains (HRgpA, RgpB and Kgp) purified from *P. gingivalis* 33277 were activated in gingipain activation buffer (20 mM HEPES, 5 mM CaCl_2_, pH 8.0; supplemented with 10 mM L-cysteine) at 37°C for 15 minutes and diluted to a final concentration of 300 nM, 75 nM and 18.5 nM. RSV and SeV particles were exposed to gingipains for 30 min at 37°C. After 30 mins, 100 μM *N*α-tosyl-L-lysine chloromethyl ketone hydrochloride (TLCK) (Millipore Sigma) was added to inhibit any residual gingipain activity. For infectivity testing, gingipain-treated viruses were added to fully confluent HEp-2 cultures, followed by 24h incubation at 37°C. Imaging of the cells was performed using the EVOS microscope. For immune blotting, gingipain-exposed viruses were precipitated by centrifugation at 25,000 x *g* for 90 min. Viruses were then resuspended in radio immunoprecipitation (RIPA) buffer with 10% 2-mercaptoethanol. Viral lysates were denatured by boiling in NuPAGE LDS Sample Buffer (Invitrogen) and NuPAGE sample reducing agent (Invitrogen) for 10 minutes, then loaded onto 4-12% gradient Bis-Tris gels (Invivogen). Proteins were then transferred into 0.2 μ PVDF membranes followed by blocking in PBS-Tween (PBST) containing 5% nonfat dried milk and incubated with primary antibodies. Anti-SeV-HN (clone M57), SeV-F (clone M16), RSV-F (Motavizumab, Thermo) and RSV-G (clone L9) were used to stain the viral glycoproteins, followed by incubation with their respective secondary antibodies. For anti-SeV-HN and SeV-F antibodies, Alexa Fluor 488 conjugated anti-mouse and anti-rabbit secondary antibodies were used, respectively. For anti-RSV-F and RSV-G, HRP-conjugated anti-human and anti-mouse secondary antibodies were used, respectively. Blots were imaged on ChemiDoc MP (BioRad).

*Histology:* HBE cultures were fixed with 4% paraformaldehyde for 15 minutes and then transferred to PBS for storage. Mouse lungs were fixed in 10% neutral buffered formalin. Tissues were submitted to the Morphology Core at the Abigail Wexner Research Center, where they were embedded in paraffin, sectioned at 5-6μm thickness, and stained with hematoxylin are eosin (H&E) or Periodic acid-Schiff (PAS) dyes.

*qPCR:* Total RNA was extracted from HBE transwells using the RNAeasy kit (Qiagen) or from mouse lungs using TRIzol reagent (Sigma-Aldrich) and converted to cDNA using the high-capacity cDNA reverse transcription kit (ThermoFisher). Proinflammatory and ISG expression was determined by TaqMan assays using TaqMan mastermix and pre-validated TaqMan probes (ThermoFisher).

*ELISA:* IFN-λ (IL-29 and IL-28), IFN-β, and CXCL10 ELISA kits were purchased from R&D Biosciences, and cytokine levels were measured in cell-free supernatants per the manufacturer’s instructions.

*RNA-seq:* Messenger RNA was purified from total RNA using poly-T oligo-attached magnetic beads. After fragmentation, the first strand cDNA was synthesized using random hexamer primers, followed by the second strand cDNA synthesis. The library was ready after end repair, A-tailing, adapter ligation, size selection, amplification, and purification. The library was checked with Qubit, and real-time PCR was used for quantification, and a bioanalyzer was used for size distribution detection. Libraries, post-QC, were pooled and sequenced on Illumina platforms according to effective library concentration and data amount. RNA-seq was done at Novogene at a depth of 50,000 paired-end reads.

*RNA-seq data analysis:* Pair-end reads from A549 cells either uninfected, challenged with *Pg* before or after IFN-lambda priming were aligned with STAR v2.7.0d (Dobin et al., 2012). Aligned reads were checked for strand specificity with infer_experiment.py script v2.6.4 from RSeQC package (Wang et al., 2012). RNA alignment metrics were assessed with CollectRnaSeqMetrics function from Picard v2.18.25 (Toolkit, 2019). Gene counts were quantified using featureCounts v2.0.0 (Liao et al., 2013) with no strand specificity parameter. Alignment and gene counts were generated against the GRCh38.p13 (GENCODE release 43) genome assembly. The DESeq2 computational pipeline v1.34.0. (Love et al., 2014) was used to normalize counts and perform differential expression analysis. Genes with less than 10 counts were filtered out. The design formula included a combination of untreated, primed, and infected conditions. DESeq analysis was performed with default parameters. For the analysis of publicly available bulk RNA-seq data, raw bulk RNA-seq counts of lung epithelial cells grown in the presence of *P. aeruginosa* PAO1 (Jonkergouw et al., 2023) were downloaded through the GEO database, accession no: GSE182847. Counts were filtered to leave only conditions of interest: negative control and PAO1 infection. The DESeq2 computational pipeline v1.34.0(Love et al., 2014) was applied as described above. The design formula included the condition of interest and replicate. Raw bulk RNA-seq counts of human nasal mucosa epithelial cells after *S. pneumoniae* challenge (Weight et al., 2019) were accessed and downloaded through phantasus web-application (Kleverov et al., 2024a) using GEO accession no: GSE124949. Counts were filtered to leave only conditions of interest: 5 days prior to the pneumococcal challenge and 2 days post for non-carrier patients. Sample S00151400 was considered an outlier and was filtered out. The DESeq2 computational pipeline v1.34.0 (Love et al., 2014) was applied as described above. The design formula included the condition of interest. Raw bulk RNA-seq counts of human lung epithelial cells infected with S. pneumoniae (Aprianto et al., 2016) were accessed and downloaded through phantasus web-application (Aprianto et al., 2016) using GEO accession no: GSE79595. Counts were filtered to leave only conditions of interest: 0, 120, and 240 minutes after infection. The DESeq2 computational pipeline v1.34.0(Love et al., 2014) was applied as described above. Timepoints 120 and 240 minutes were combined as late time points of infection. The design formula included timepoint and pneumococcal strain type. For downstream enrichment analysis, only results coming from the wild-type strain were used.

*Enrichment analysis* and *Visualization of transcriptional data:* Gene set enrichment analysis was performed on the pre-ranked list of genes using the R package fgsea v1.27.1 (Korotkevich et al., 2021). Data ranking was performed based on Wald statistics. Pathway enrichment was performed using canonical pathway database collection accessed through msigdb R package v7.5.1(Dorgalev, 2022). PCA plot was created using plotPCA function from DESeq2 v1.34.0 (Love et al., 2014) after regularized log transformation. Bar plots were created using ggplot2 package v3.4.4.9000 (Wickham, 2016). Heatmaps were visualized using the phantasus web-application (Kleverov et al., 2024b).

*Flow cytometric measurement of RSV replication:* HEp-2 cells seeded in 12-well plates were infected with *P. gingivalis* (MOI 100) for 4h at 37°C, followed by RSV infection (1.2x10^4^PFU) for 2h. After infection cells were lifted using trypsin EDTA (0.05%), then washed to remove the trypsin and analyzed by flow cytometry using a LSR Fortessa X-20 cytometer (BD).
