## Supplementary figures and images for "Virion stripping: A new role for bacterial proteases"

### Supplemental Figure S1

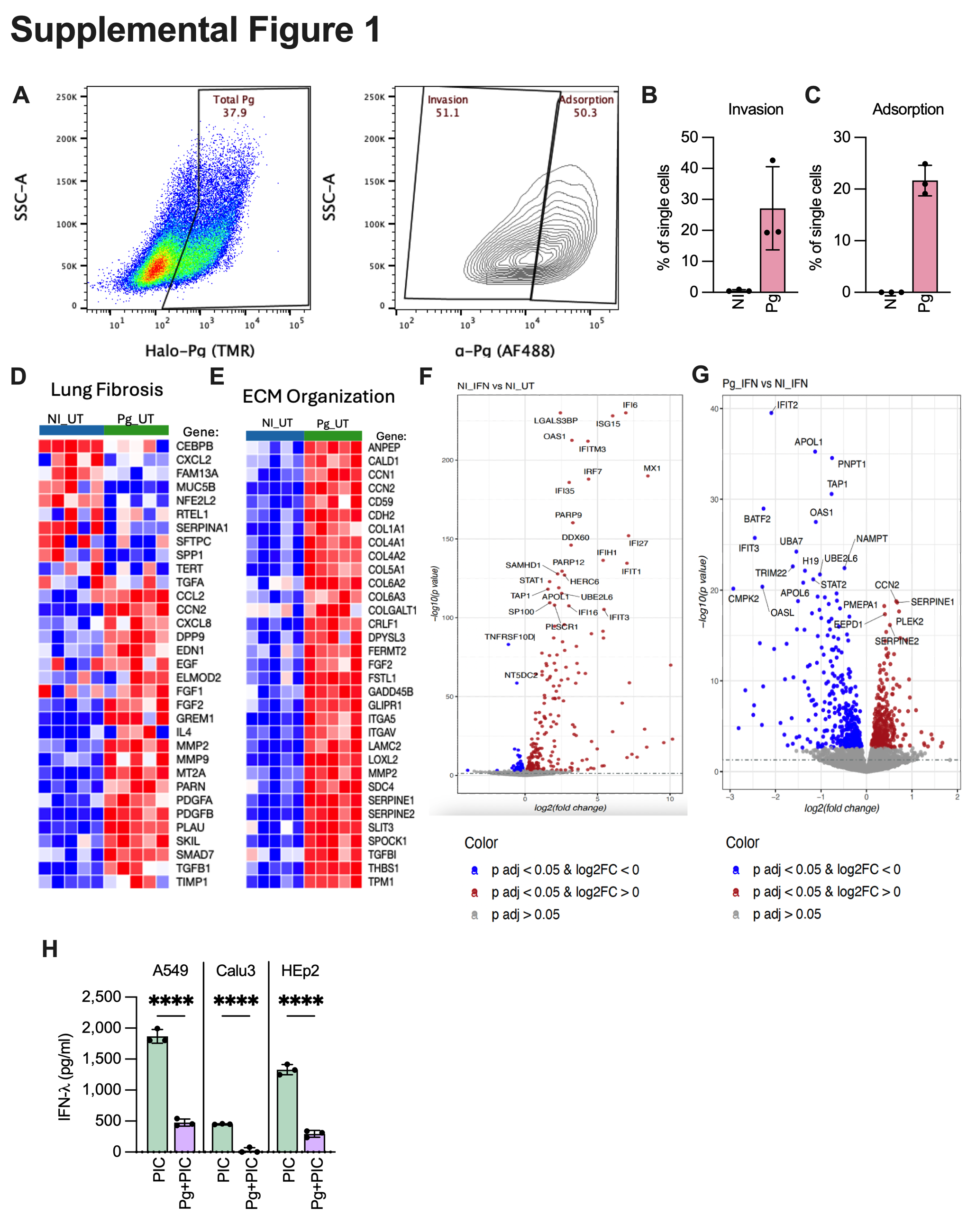

### Supplemental Figure S2

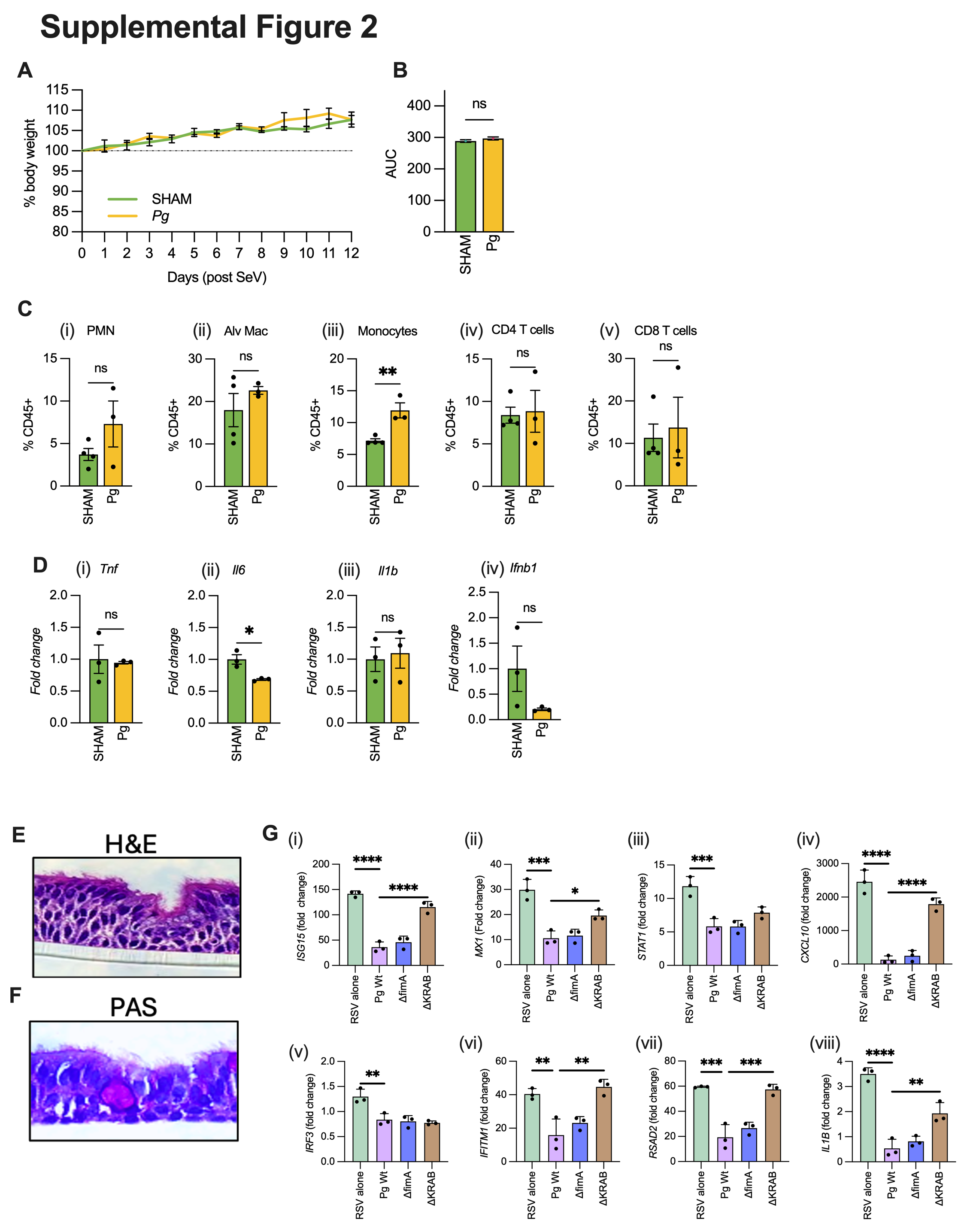

### Supplemental Figure S3

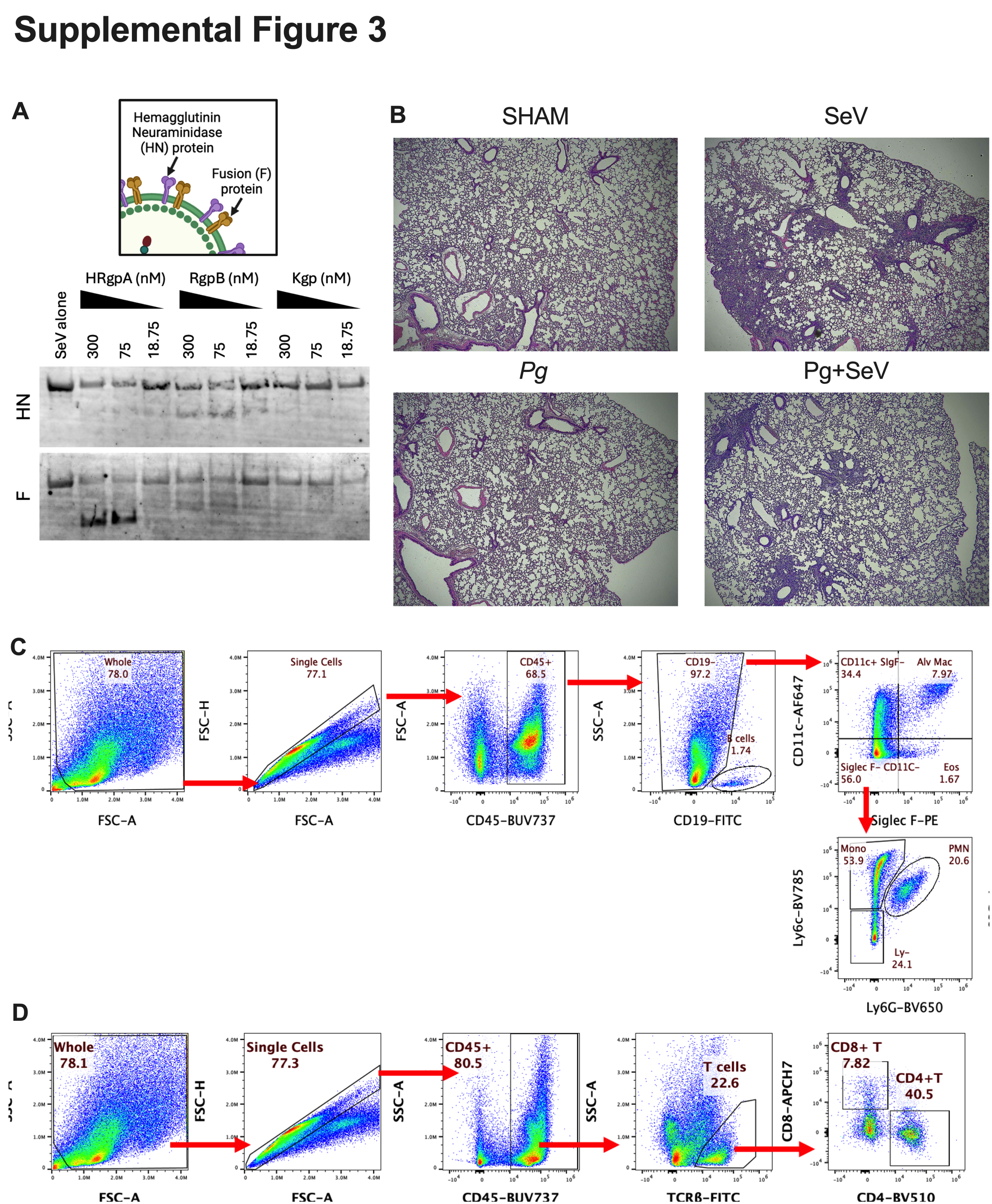
